# Syndecan-3 enhances anabolic bone formation through WNT signalling

**DOI:** 10.1101/846972

**Authors:** Francesca Manuela Johnson de Sousa Brito, Andrew Butcher, Addolorata Pisconti, Blandine Poulet, Amanda Prior, Gemma Charlesworth, Catherine Sperinck, Michele Scotto di Mase, George Bou-Gharios, Robert Jurgen van ’t Hof, Anna Daroszewska

**Affiliations:** Department of Musculoskeletal Biology I, Institute of Ageing and Chronic Disease, University of Liverpool, L7 8TX, UK; Department of Biochemistry, IIB, University of Liverpool, L7 8TX, UK; Department of Biochemistry and Cell Biology, Stony Brook University, Stony Brook, NY 11794-5215, USA; Department of Clinical Biochemistry and Metabolic Medicine, The Royal and Broadgreen University Hospitals NHS Trust, Liverpool, L7 8XP, UK; Department of Rheumatology, The Royal and Broadgreen University Hospitals NHS Trust, Liverpool, L7 8XP, UK

**Keywords:** Syndecan-3, bone, osteoblast, osteoclast, WNT, Frizzled

## Abstract

Osteoporosis is the most common age-related metabolic bone disorder, which is characterised by low bone mass and deterioration in bone architecture, with a propensity to fragility fractures. The best treatment for osteoporosis relies on stimulation of osteoblasts to form new bone and restore bone structure, however anabolic therapeutics are few and their use is time-restricted. Here we report that Syndecan-3 (SDC3) increases new bone formation through enhancement of WNT signalling. Young adult *Sdc3^−/−^* mice have a low bone volume phenotype associated with reduced bone formation, increased bone marrow adipose tissue (BMAT), increased bone fragility and a blunted anabolic bone formation response to mechanical loading. The premature osteoporosis-like phenotype of *Sdc3^−/−^* mice is primarily explained by delayed osteoblast maturation and impaired osteoblast function, with contributing increased osteoclast-mediated bone resorption. Mechanistically, SDC3 enhances canonical WNT signalling in osteoblasts through stabilisation of Frizzled 1, making SDC3 an attractive target for novel anabolic drug development.

## INTRODUCTION

Osteoporosis is the most common age-related metabolic bone disorder affecting millions of people worldwide and predicted to rise as the population ages^1^. Osteoporosis is characterised by low bone mass and impaired bone micro-architecture, which predispose to fragility fractures, approximately 9 million annually^2^. Only a small number of anabolic treatments is available, use of which is time-limited and restricted to severe cases^3^. As osteogenic therapeutics are better than anti-resorptives, due to their ability to restore bone mass and architecture, there is a pressing need to gain a better understanding of anabolic pathways leading to bone formation so that novel management strategies can be developed.

One of the most important signalling pathways inducing osteoblastogenesis and bone formation is WNT-dependent^4^. Mechanical loading of bone induces osteogenesis through the activation of the canonical (β-catenin-dependent) WNT-signalling, subsequent to repression of sclerostin, a WNT inhibitor expressed by mechanosensing osteocytes.^5^

The canonical WNT-signalling pathway is activated by WNT ligands, which bind to the receptor complex comprised of frizzled (FZD) and the low density lipoprotein receptor-related proteins 5/6 (LRP5/6) to indirectly inhibit phosphorylation of β-catenin, allow for its translocation into the nucleus and induction of β-catenin-response genes including *AXIN2*^4^. WNT signalling intensity depends on the availability of FZD receptors, the level of which is regulated by ubiquitin-mediated degradation involving E3 ubiquitin ligases (ZNRF3 or RNF43), dependent on, or independent of leucine-rich repeat containing G protein-coupled receptors (LGRs)^6,7^. In the presence of extracellular R-spondins (RSPOs), which bind to LGRs, the RSPO-LGR complex interacts with ZNRF3/RNF43 and prevents the latter from tagging FZD for degradation. Thus, the resulting high level of FZD allows for strong WNT-induced signalling^6^. Inactivating mutations of RSPO2, which interfere with RSPO2-LGR or - RNF43 interaction, inhibit WNT signalling and induce limb abnormalities in humans^7^ and mice^8^. However, as deletion of LGRs does not recapitulate this phenotype, another, LGR-independent, mechanism of regulation of FZD level has been proposed^7^ and evidence presented indicating that it may involve heparan sulphate proteoglycans (HSPGs), including syndecans (SDCs)^6,9^. Interestingly, a gene expression study of human bone subject to mechanical loading revealed that some of the induced pathways are mediated by SDCs^10^.

SDCs are a family of four type 1 transmembrane HSPG receptors, which play a role in cell adhesion, signalling, and are co-receptors for growth factors and their receptors. SDCs contain three domains: an N-terminal ectodomain, a transmembrane- and a C-terminal cytoplasmic domain. The ectodomain, which can be shed, mediates cell-cell and cell-matrix interaction through the attached glycosaminoglycan (GAG) chains including heparan sulfate (HS) in case of SDC2 and SDC4, and both HS and chondroitin sulfate (CS) in case of SDC1 and SDC3. The transmembrane domain allows for dimerization, whereas the cytoplasmic domain (composed of conserved regions C1 and C2 flanking a variable region [V]) interacts with kinases, the cytoskeleton and C1 mediates endocytosis^11,12^.

GAGs, abundant in the extracellular matrix (ECM), are known to enhance bone regeneration and osteoblastogenesis by binding of the WNT inhibitor, sclerostin. However, with ageing the amount of GAGs, and in particular CS decreases in human cortical mineralised bone matrix, which has a detrimental effect on bone toughness and likely contributes to the age-related deterioration of bone quality^13^. Previous research^14^ has shown that HS GAGs are important in osteoblast differentiation and activity, and SDCs with attached GAGs, can modulate WNT signalling^15,16^, however SDC-WNT crosstalk in bone has not been clearly deciphered^4,17^ and involvement of SDCs in bone physiology is poorly defined. SDC3 is expressed in the bone shafts of developing chick embryos^18^ and plays a complex role in regulating chondrocyte proliferation^19,20^, however its exact role in skeletogenesis or bone homeostasis is not fully understood.

Here we demonstrate that deletion of *Sdc3* leads to a premature low bone volume phenotype associated with increased bone fragility and increased bone marrow adipose tissue (BMAT) in young adult mice. The underlying mechanism involves impaired bone formation and attenuated anabolic bone response to mechanical loading, explained by impaired canonical WNT3a-mediated signalling and likely due to increased degradation of FZD1 receptor in osteoblast-lineage cells.

## MATERIALS AND METHODS

### Mice

*Sdc3^−/−^* mice were housed in a pathogen-free facility at the University of Liverpool, with free access to food and water, and in accordance with the Animals (Scientific Procedures) Act 1986 and the EU Directive 2010/63/EU, after the local ethical review and approval by Liverpool University’s Animal Welfare and Ethical Review Body (AWERB). *Sdc3^−/−^* mice on C57Bl6 background^21^ were donated by Dr Heikki Rauvala, University of Helsinki, Finland. Sdc3^−/−^ mice were generated from heterozygous and/or one generation homozygous breeding. Control wild type (WT) mice were littermates or closely related, age- and sex-matched.

### *In vivo* mechanical loading

The right tibiae of 10-week-old male *Sdc3^−/−^* (n=8) and WT (n=7) mice were mechanically loaded to induce metaphyseal bone formation, as described^22^. Briefly, mice were anaesthetised using Isoflurane through the whole procedure. The right knee and ankle were placed in custom made cups in the ElectroForce® 3100 Test Instrument (Bose Corporation, MN, USA) and repetitive mechanical compression applied at a peak magnitude of 11N for 0.05sec. Baseline hold time of 2N was applied for 9.9sec to maintain the tibia in place in between peak load applications. Each loading episode consisted of 40 cycles and was applied 6 times over a period of 2 weeks. Mice received calcein (2mg/ml) 150µl intraperitoneal injections on days 8 and 12 (day 1 = first loading day). Mice were culled 3 days after the final loading episode and tibiae collected.

### µCT analysis

The skin was removed, and hind limbs and spines were fixed overnight in 4% formalin-buffered saline and stored in 70% ethanol. µCT analysis was performed using a Skyscan 1272 system. For assessment of bone morphometry, tibias, femurs and the lumbar spine were dissected free of most soft tissue and scanned at a resolution of 4.5 µm (60 kV, 150 µA, rotation step size 0.3°, using a 0.5 mm aluminium filter). The reconstruction was performed using the Skyscan NRecon package. Trabecular parameters were measured using Skyscan CTAn software in a stack of 200 slices immediately proximal, or distal to the growth plate in femurs, or tibias respectively, as described previously^23^. Tibial and femoral cortical parameters were measured using Skyscan CTAn software in a stack of 100 slices immediately proximal to the tibio-fibular junction and distal to the trochanter respectively.

Trabecular bone was separated from cortical bone using a CTAn macro, and the same macro then performed measurement of the cortical and trabecular parameters according to ASBMR guidelines^24^. For tibial length quantification, the full length of mineralised tibial diaphysis at P2, and the distance between proximal and distal growth plate during skeletal maturation was measured on µCT longitudinal scans of 9 µm resolution using the Data Viewer.

For measurement of the whole bone volume of the tibial bone shaft of 2 day-old and 2-week old mice, tibias were scanned at a resolution of 9µm, and reconstructed as detailed above. Using a macro in CTAn, the bone was separated from soft tissues using a fixed threshold and bone volume measured between the proximal and distal growth plates.

### Bone histomorphometry and histology

Bone samples were processed and stained for histology as previously described^25^. Briefly, mice received intraperitoneal calcein injections (2mg/ml, 150µl) 5 days and 2 days before culling. The skin was removed, hind limbs were fixed for 24h in 4% formalin, and stored in 70% ethanol. The samples were embedded in methyl methacrylate (MMA) and 5µm sections were cut using a tungsten steel knife on a Leica motorised rotary microtome. Sections were stained for TRAcP to visualise osteoclasts and counterstained with Aniline Blue. For analysis of calcein double labelling, sections were counterstained with Calcein Blue and histomorphometry performed as described previously^25^. Sections were visualised on a Zeiss Axio Scan.Z1 using a x10 lens (pixel size 0.45 µm) for TRAcP stained sections, and a x20 lens and a pixel size of 0.23 µm for calcein labels. Histomorphometry was performed using the TrapHisto, OsteoidHisto and CalceinHisto open-source image analysis programmes^25^ available at https://www.liverpool.ac.uk/ageing-and-chronic-disease/bone-hist/.

Adipocytes were quantified in sections stained with Goldner’s Trichrome, imaged on the Zeiss Axio Scan.Z1 using a x10 lens, and quantified using an in-house developed program (FatHisto) based on ImageJ. The presence of adipocytes in the MMA-embedded sections was verified by immunostaining for perilipin on dewaxed paraffin-embedded sections of tibiae from 3-month old *Sdc3^−/−^* (n=3) and WT (n=3) mice. Sections were incubated in Unitrieve solution, blocked with goat serum and adipocytes were detected using perilipin antibody (Abcam) and secondary goat anti-rabbit Alexa Fluor 594 antibody (Invitrogen) at 1:500 and 1:200 dilution respectively.

For analysis of the growth plate, mouse knees were decalcified in formical overnight, embedded in wax and 5 µm sections cut on a Leica rotary microtome. Sections were stained for haematoxylin as well as Safranin O/fast Green, imaged using the Zeiss Axio Scan.Z1 using a x10 lens, and growth plate width measured using Zeiss Zen software.

### Three-point bending test

Flexural strength of the femurs from 3-month old *Sdc3^−/−^* and WT mice was assessed by dynamic three-point bending test on freshly dissected, unfixed bones using the ZWICK-mechanical testing machine (ZWICK, Switzerland), using a span of 8mm between the support points and a crosshead speed of 1 mm/min.

### Osteoclast culture

Bone marrow was obtained from 2-3-month-old mice and cultured with 100 ng/ml of M-CSF (Prospec Bio) in αMEM (Gibco Invitrogen) supplemented with 10% FCS for three days. Next the adherent M-CSF dependent macrophages were harvested and plated in 96-well plates at 10^4^ cells/well in αMEM supplemented with 10% FCS, 25 ng/ml M-CSF and varying amounts of RANKL (R&D Systems), according to standard methods as previously described^26^. After 5-6 days of culture, the cells were fixed using 4% paraformaldehyde in PBS, washed and stained for TRAcP. Numbers of osteoclasts were counted after TRAcP staining by an observer blinded to the genotype. Resorption assays were performed using the same method, but by plating the cells on dentine slices placed within 96-well plates. After six days, the plates were fixed and stained for TRAcP, and number of osteoclasts per dentine slice counted. Next the cell layer was polished off the dentine slices, and the entire slices imaged using an Olympus reflected light microscope (5x lens) fitted with a Zeiss axiocam camera (isotropic pixel size 1.2µm). Individual fields (8-9 per slice) were stitched together using Microsoft ICE, and the pit area measured using ImageJ.

### Osteoblast culture

Bone marrow was obtained from the long bones of 2-3-month-old mice by removing the ends of the bones to expose the bone marrow cavity, followed by centrifugation at 300g for 3 min. Bone marrow mesenchymal stromal cells (BMSCs) were cultured in DMEM (Gibco Invitrogen) supplemented with 10% FCS and antibiotics (non-osteogenic culture conditions). Osteoblasts were differentiated from BMSCs by culturing in osteogenic medium, DMEM supplemented with 10% FCS, 1% P/S, 2mM β-glycerophosphate (Sigma-Aldrich) and 50µg/ml 2-Phospho-L-ascorbic acid trisodium salt (Sigma-Aldrich). After removal of the bone marrow, the remaining femoral and tibial diaphysis were cut into small pieces followed by collagenase digestion for 30 min. The bone chips were washed in PBS and cultured in αMEM supplemented with 10% FCS in 25 cm^2^ tissue culture flasks until semi confluent layers of osteoblasts (bone chip osteoblasts, BC) had grown out of the bone chips. The cultures were washed with PBS and the osteoblasts harvested using trypsin.

Calvarial osteoblasts were isolated from the calvaria of 2-3 day-old pups by collagenase digestion as described in detail previously^27^ and cultured in αMEM supplemented with 10% FCS and antibiotics.

For alkaline phosphatase assays, BC osteoblasts were plated at 15×10^3^ cells/well in 96-well plates and cultured in 150 µl αMEM with 10% FCS for 24 hours. Alamar Blue (15 µl/well) was added to the culture and culture continued for 2 more hours. Cell viability was measured by analysing the Alamar Blue signal using a fluorescence plate reader (BMG FLUO Star Optima, excitation 560 nm, emission 590 nm). Next the cells were fixed for 10 min at 4°C using 4% paraformaldehyde in PBS, washed and stored dry at −20 °C. Alkaline phosphatase was measured using the conversion of para-nitrophenyl-phosphate by measuring absorption at 405 nm at 1 min intervals for 30 min. Alkaline phosphatase measurements were corrected for cell number by dividing by the Alamar Blue signal.

For mineralisation assays osteoblasts, either derived from bone chips or calvaria, were cultured for 26 days in mineralising-medium containing DMEM (Gibco Invitrogen) supplemented with 10% FCS, 1% P/S, 2mM β-glycerophosphate (Sigma-Aldrich) and 50µg/ml 2-Phospho-L-ascorbic acid trisodium salt (Sigma-Aldrich). Half medium changes were made every other day during induction period and mineralising-medium was made up fresh every time. Calcium deposition in the mineralisation assay, at day 26, was measured using Alizarin Red staining. Osteoblasts were fixed in 70% ice cold Ethanol for 1h on ice and washed in dH_2_O. Cells were then incubated in Alizarin Red S (Sigma-Aldrich) solution for 20 minutes at room temperature on a rocking table, washed in dH_2_O overnight and then air dried. Quantification was performed by dissolving the alizarin Red in 10% (w/v) cetylpyridinium chloride in 10mM sodium phosphate, and measuring absorbance at 562 nm as previously described^28^. For mineralisation assays of BMSC, cells were grown to confluence in 12 well plates in DMEM supplemented with 10% FCS and antibiotics. Medium was changed to osteogenic medium and culture continued for 10 days with half medium changes every other day. Cultures were fixed and stained for Alizarin Red S as described above.

### Adipo-osteogenic culture of bone marrow stromal cells

BMSCs were isolated from mouse long bones as described above, plated at 2×10^6^ cells per cm^2^ in αMEM supplemented with 15% FCS and grown until confluent (approximately 10 days) with twice weekly medium changes. After reaching confluence the medium was replaced with co-differentiation adipogenic and osteogenic medium, αMEM supplemented with 10% FCS, 2mM β-glycerophosphate, 50µg/ml 2-Phospho-L-ascorbic acid trisodium salt and 100 nM dexamethasone (Sigma-Aldrich), adapted from Ghali *et al*^29^. Culture was continued with 3-weekly changes of medium for 12 more days, plates fixed, and adipocytes stained with Oil-red-O. After staining, the plates were imaged using a Nikon Diaphot inverted microscope fitted with a Touptec camera, using a 10x lens, and number of adipocytes per image field counted, counting at least 3 fields for every mouse cell donor.

### Cell proliferation assay

Anti-Ki67 (Abcam) was used to determine cell proliferation. BC-differentiated osteoblasts from Sdc3^−/−^ (n=6) and WT (n=7) mice were plated on glass cover slips (6 per mouse) in 24-well plates, cultured for 48h, fixed in 100% Methanol for 5 minutes, permeabilized with 0.1% Triton-X-100 for 5 minutes and then blocked with 10% FCS in PBS for 1h. The cells were then incubated with Anti-Ki67 antibody at 1:250 dilution overnight at 4°C. Fluorescent Alexa-fluor-488 conjugated antibody (Life Technologies) at 2µg/ml was used for detection. Nuclei were stained with DAPI. The coverslips were imaged using a Zeiss Axio Scan.Z1 using a x20 lens, and Ki-67 positive and negative nuclei quantified using a macro in ImageJ.

### qPCR

RNA was extracted from cell cultures using Trizol. Reverse transcription was performed (1 µg RNA/sample) with 3 replicate RT reactions per sample, using Roche EvoScript according to the manufacturers protocol. qPCR was performed on a Roche Lightcycler 480 using Roche Lightcycler mastermix and validated Roche and Fisher Scientific TaqMan primer probe sets (Supplementary Table 1). *Hmbs* was used as a house keeping gene, and expression was calculated as relative expression to this house keeping gene using the delta-CT method. For experiments on the effects of WNT3a, cells were cultured as described until near-confluence and after overnight serum starvation (where 10% FCS was replaced by 2% TCM), recombinant mouse WNT3a protein (Abcam) 50ng/ml was added and cells cultured for 8 hours before lysis in Trizol.

### Western Blotting

Cells were cultured as described above. After cell lysis using standard RIPA buffer, proteins were separated by sodium dodecyl sulphate-polyacrylamide gel electrophoresis and electroblotted onto nitrocellulose membranes (Bio-Rad). Membranes were blocked with blocking buffer (LI-COR) and probed with primary antibodies to FZD1 (1:750, R&D), β-catenin and GAPDH (1:1000, Cell Signalling). After washing with TBST, membranes were incubated with appropriate LI-COR infrared-labelled secondary antibodies, visualised using the Odyssey CLx imaging system (LI-COR) and analysed using Image Studio lite (LI-COR, 5.2).

### Statistical analysis

All experiments were performed with a minimum of 3 biological replicates, unless indicated otherwise. Statistical analysis was performed in SPSS using a (two-tailed) Student’s *t* test, unless indicated otherwise and a *p* < 0.05 was considered statistically significant.

## RESULTS

### Syndecan-3 deletion results in a low bone volume phenotype in adult mice

Morphometric µCT analysis of proximal tibiae of 3 month-old male *Sdc3^−/−^* mice revealed a decrease in bone volume (BV/TV), trabecular thickness (Tb.Th) and trabecular number (Tb.N) of 34% (*p<0.001*), 13% (*p<0.001*) and 24% (*p<0.001*) respectively and an increase in trabecular separation (Tb.Sp), trabecular pattern factor (Tb.Pf, indicating reduced connectivity), and structure model index (SMI, indicating a more rod-like rather than plate-like structure), of 10% (*p<0.05*), 50% (*p<0.001*) and 20% (*p<0.001*) respectively compared to WT mice (Fig. 1a,b and Supplementary Table 2a). Morphometry of distal femora showed similar results in keeping with a low trabecular bone volume phenotype in young adult *Sdc3^−/−^* mice (Fig. 1b and Supplementary Table 2a).

**Fig. 1.**
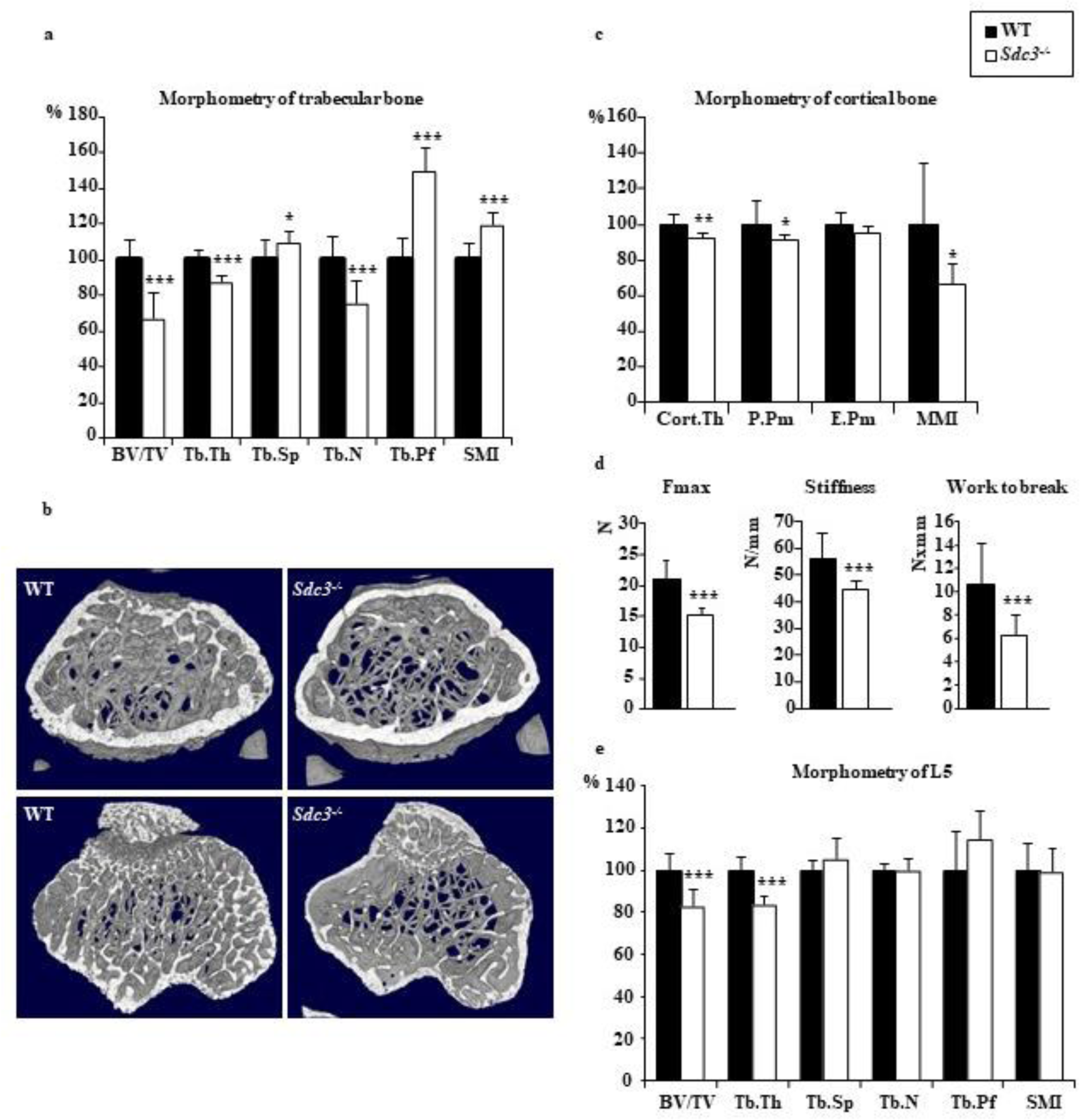
Syndecan-3 enhances bone volume, architecture and strength in adult mice. **(a)** Morphometry of proximal tibiae of 3-month old male *Sdc3^−/−^* (n=13) and WT (n=12) mice analysed using µCT. BV/TV: bone volume, Tb.Th: trabecular thickness, Tb.Sp: trabecular separation, Tb.N: trabecular number, Tb.Pf: trabecular pattern factor, SMI: structure model index. **(b)** Crossection image of a 3D µCT reconstruction of distal femurs (top panels) and proximal tibiae (bottom panels) of *Sdc3^−/−^* vs WT mice representing the volumes used for analysis of trabecular bone parameters. µCT at 4.5 µm. **(c)** Morphometry of tibial cortex of 3-month old male *Sdc3^−/−^* (n=10) and WT (n=12) mice analysed using µCT. C.Th: cortical thickness, E.Pm: endosteal perimeter, P.Pm: periosteal perimeter, MMI: polar mean moment of inertia. **(d)** Three-point bending test on femurs assessing the maximum force (Fmax), stiffness and work to break in *Sdc3^−/−^* (n=10) vs WT (n=10) male mice. **(e)** Morphometry of spine (L5) of 3-month old male *Sdc3^−/−^* (n=11) and WT (n=11) mice. Data in a and c-e are shown as mean ± SD. * *p<0.05*, ** *p<0.01*, *** *p<0.001*

Cortical thickness (C.Th), periosteal perimeter (P.Pm) and polar mean moment of inertia (MMI) were decreased by 8% (*p<0.01*), 9% (*p<0.05*) and 34% (*p<0.05*) respectively in the tibiae of 3-month old male *Sdc3^−/−^* mice vs WT (Fig. 1c and Supplementary Table 2b). Morphometry of femoral cortex showed a decrease in C.Th, P.Pm, endosteal perimeter (E.Pm) and MMI of 11% (*p<0.001*), 6% (*p<0.01*), 12% (*p<0.001*) and 21% (*p<0.01*) respectively in *Sdc3^−/−^* vs WT (Supplementary Table 2b). As expected from the results of the µCT analysis, the three-point bending test applied to femurs revealed a 27% (*p<0.001*) and 40% (*p<0.001*) reduction in maximum force and work to fracture respectively in the *Sdc3^−/−^* vs WT mice, as well as a 20% reduction in stiffness (*p<0.001*) (Fig. 1d), indicative of bone fragility in *Sdc3^−/−^* mice. Spinal morphometry showed a decrease in BV/TV and Tb.Th in *Sdc3^−/−^* vs WT as well (Fig. 1e). Thus, *Sdc3* deletion induces a low bone volume phenotype in both the trabecular and cortical compartments in adult mice, which is associated with bone fragility.

### Syndecan-3 differentially regulates prenatal and postnatal bone development

To ascertain whether the low bone volume phenotype of adult *Sdc3*^−/−^ mice was due to a developmental defect, we measured the total bone volume and length of tibiae just after birth and during skeletal maturation. Surprisingly, tibiae from 2-day old *Sdc3^−/−^* pups had an approximately 40% higher bone volume vs WT (*p<0.001*), however by 2 weeks of age the *Sdc3^−/−^* mice tibia bone volume was approximately 20% lower than that of the WT (*p<0.01*; Fig. 2a). There was no difference in trabecular BV/TV between the genotypes at 6 weeks, however as of 8 weeks of age the BV/TV was significantly lower in *Sdc3^−/−^* vs WT mice (*p<0.001*; Fig. 2b). This differential bone volume variation was mirrored by changes in the tibial longitudinal growth. Thus at 2 days of age the tibiae of *Sdc3^−/−^* mice were 0.8 mm longer than those of WT (*p<0.001*; Fig. 2c), however this difference was reversed already by 2 weeks, at which stage the *Sdc3^−/−^* tibiae were shorter by 0.9 mm vs WT (*p<0.001*) and remained shorter throughout (Fig.2c). Taken together this data indicate that SDC3 has a differential effect on pre- and post-natal longitudinal bone development and growth.

**Fig. 2.**
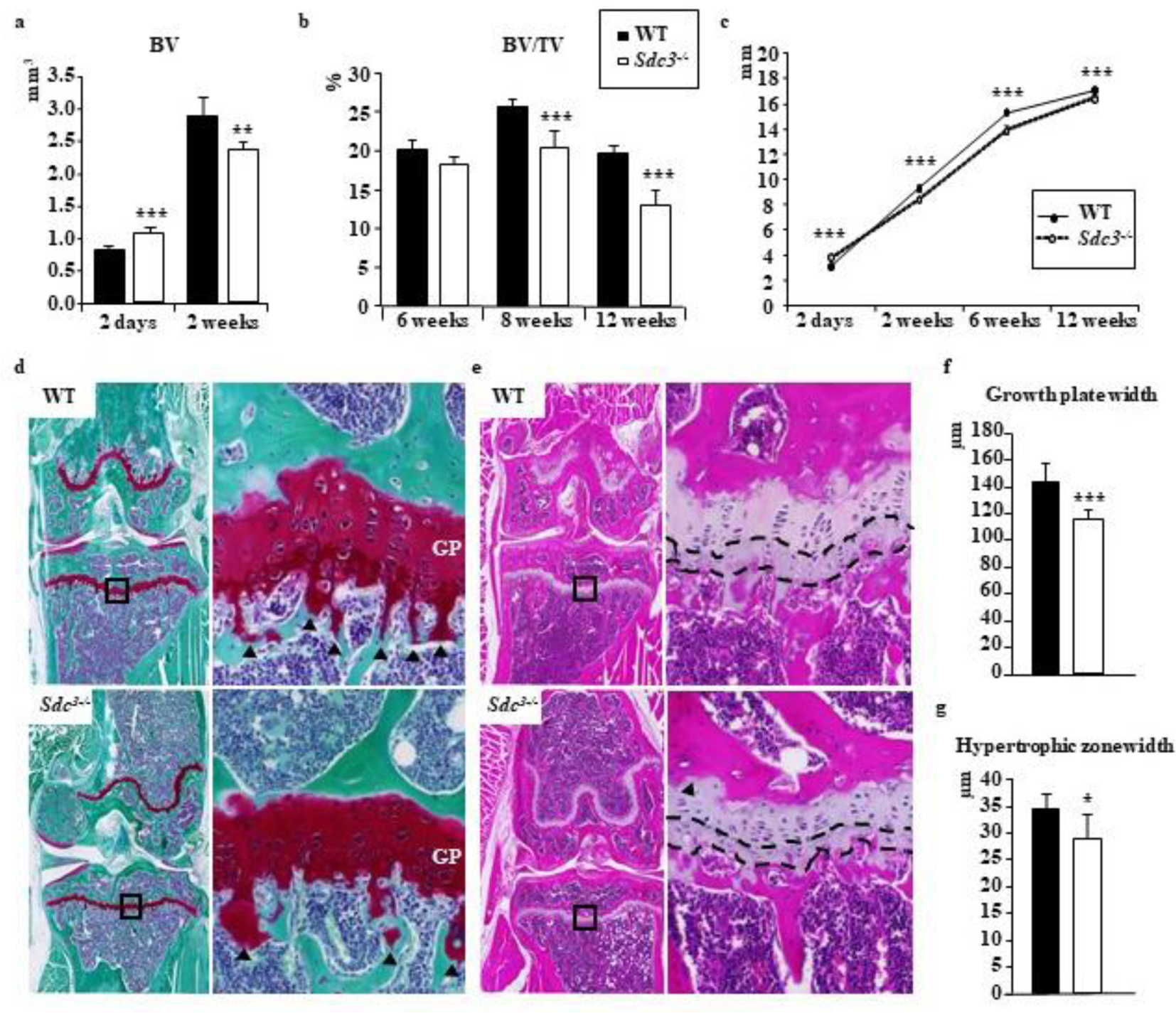
Syndecan-3 differentially affects bone development *in utero* vs postnatally. **(a)** Total bone volume of tibiae from 2-day old pups (*Sdc3^−/−^* n=9 vs WT n=9) and 2-week old male mice (*Sdc3^−/−^* n=6 vs and WT n=6) quantified using µCT. **(b)** Trabecular bone volume per tissue volume (BV/TV) of the proximal tibia in males assessed by µCT during skeletal maturation at weeks: 6, 8 (at each time-point *Sdc3^−/−^* n=6 vs WT n=6) and 12 (*Sdc3^−/−^* n=13 vs WT n=11). **(c)** Length of tibial diaphysis quantified on µCT during skeletal maturation after birth at day 2 (*Sdc3^−/−^* n=9 vs WT n=9), week 2 (*Sdc3^−/−^* n=6 vs WT n=6), 6 (*Sdc3^−/−^* n=5 vs WT n=6) and 12 (*Sdc3^−/−^* n=6 vs WT n=5). All mice were male except pups at day 2 (undefined). **(d)** Safranin-O/fast green stain of knee sections of 3-month old WT and *Sdc3^−/−^* mice. Regions of the growth plate delineated by the squares in the left panels have been magnified in the right panels. Primary spongiosa is indicated by arrowheads. GP - growth plate. **(e)** H&E stain of knee sections of 3-month old WT and *Sdc3^−/−^* mice. Regions of the growth plate delineated by the squares in the left panels have been magnified in the right panels. Hypertrophic region of the growth plate is indicated by dashed lines. Disorganised morphology of the growth plate in *Sdc3^−/−^* is indicated by an arrowhead. **(f)** Quantification of the growth plate width in knee sections of 3-month old male WT (n=8) and *Sdc3^−/−^* (n=4) mice. **(g)** Quantification of the growth plate hypertrophic zone in knee sections of 3-month old male WT (n=8) and *Sdc3^−/−^* mice (n=4).Data in a, b, c, f and g are shown as mean ± SD. \**p<0.05*, \*\**p<0.01*, \*\*\**p<0.001*

As SDC3 is known to attenuate BMP2-mediated limb chondrogenesis^30^, we measured the growth plate width in the proximal tibia of 3-month old mice, and found it thinner in *Sdc3^−/−^* vs WT mice, *p<0.001* (Fig. 2d and f). Furthermore, the hypertrophic zone width was reduced, *p<0.05* (Fig. 2e and g), and disorganised growth plate morphology was evident, with poorly stacked chondrocytes in irregular columns within the proliferating zone and a virtual absence of primary spongiosa (Fig.2d and e), which in part may explain the decreased long bone growth in young and adult *Sdc3^−/−^* mice.

### Low bone volume phenotype in adult *Sdc3^−/−^* mice is associated with impaired osteoclastogenesis, but increased osteoclast resorption of bone

Bone histomorphometry of 3 month-old *Sdc3^−/−^* mice after TRAcP staining for osteoclasts unexpectedly revealed a decrease of 44% in both osteoclast surface and number per bone surface (Oc.S/BS, *p<0.05* and N.Oc/BS, *p<0.01* respectively) vs WT. Likewise, the number of osteoclasts per tissue volume (N.OC/TV) was decreased by 57% in *Sdc3^−/−^* vs WT (*p<0.01*; Fig.3a and b, and Supplementary Table 3).

**Fig.3.**
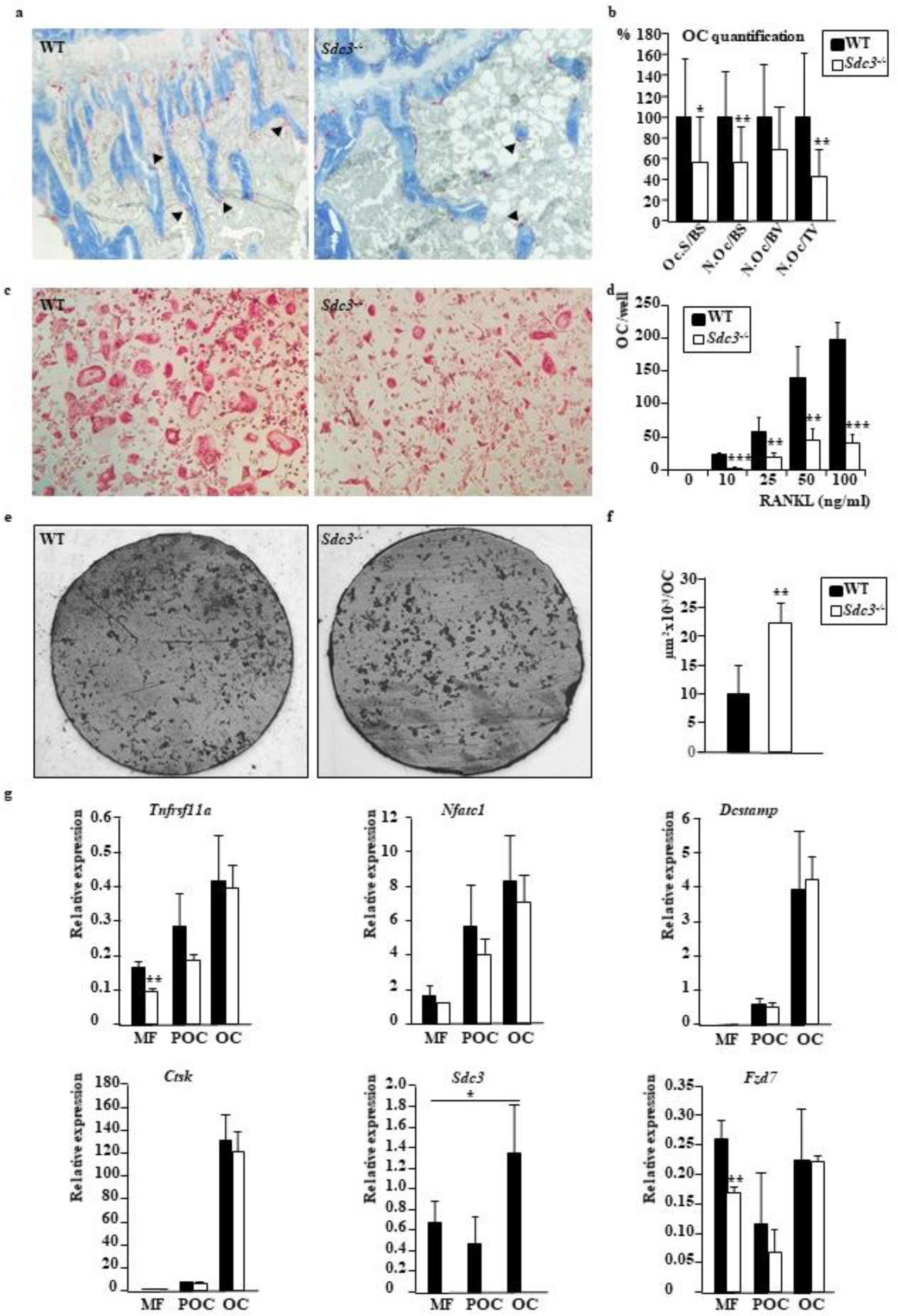
Syndecan-3 enhances osteoclastogenesis but modulates osteoclastic bone resorption. **(a)** Histology of proximal tibia showing osteoclasts TRAcP-stained red (indicated by arrow-heads), bone counterstained with aniline blue of 3-month old WT (left panel) and *Sdc3^−/−^* (right panel) mice.**(b)** Quantification of TRAcP-stained osteoclasts on histology of tibiae from 3-month old WT (n=12) and Sdc3^−/−^ (n=12) mice. Oc.S/BS: osteoclast surface/bone surface, N.Oc/BS: number of osteoclasts/bone surface, N.Oc/BV: number of osteoclasts/bone volume, N.Oc/TV: number of osteoclasts/tissue volume. OC: osteoclast. **(c)** Photomicrographs of TRAcP-stained osteoclasts differentiated *in vitro* from RANKL- and MCSF-stimulated macrophages of WT (left panel) and *Sdc3^−/−^* (right panel) mice. **(d)** Quantification of osteoclast numbers in RANKL- and MCSF-stimulated macrophage cultures from WT and *Sdc3^−/−^* mice. RANKL stimulation was at 10, 25, 50 and 100 ng/ml. Data shown are from one representative experiment out of 5 performed (with 5 replicate wells per mouse cell donor). **(e)** Reflected light microphotographs of dentine slices with areas (in black) resorbed by osteoclasts differentiated from WT (left panel) and *Sdc3^−/−^* (right panel) mice. **(f)** Quantification of resorbed area on dentine slices by osteoclasts from WT and *Sdc3^−/−^* mice (n=3; 4 dentine slices per mouse). **(g)** Relative expression of *Tnfrsf11a, Nfatc1, Dcstamp, Ctsk, Sdc3* and *Fzd7* to *Hmbs* in MCSF-dependent macrophages (MF), pre-osteoclasts (POC) and osteoclasts (OC) differentiated from *Sdc3^−/−^* (n=3) and WT (n=4) mice. qPCR reactions were performed in triplicate. There was a statistically significant increase in expression of *Tnfrsf11a* (*p<0.05*), *Dcstamp* (*p<0.01*), *Nfatc1* (*p<0.01*) and Ctsk (*p<0.001*) between the MF and OC stage in both *Sdc3^−/−^* and WT (not annotated for clarity). Data in b, d, f and g are shown as mean±SD, * *p<0.05*, ** *p<0.01*, *** *p<0.001*

The rate of osteoclast formation induced by MCSF and RANKL *in vitro* was significantly lower in osteoclastogenesis cultures derived from *Sdc3^−/−^* vs WT 3-month old mice (Fig.3c and d), confirming the histology findings (Fig.3a and b), and indicating that the decreased osteoclast formation is due to a cell autonomous defect in response to RANKL. Interestingly, cultures of osteoclasts on dentine slices did not show a difference in resorption area between *Sdc3^−/−^* and WT (Fig.3e), however due to there being fewer *Sdc3^−/−^* osteoclasts vs WT formed per dentine slice (80.6±18.4 vs 141.2±98.5 respectively, *p<0.05*), the resorption area per osteoclast was over twofold higher in *Sdc3^−/−^* vs WT cultures, *p<0.01* (Fig.3f). These findings suggest that SDC3 promotes osteoclast differentiation but decreases mature osteoclast resorptive activity.

To gain insight into the mechanistic role of SDC3 in osteoclastogenesis, we examined osteoclast marker gene expression during osteoclast differentiation *in vitro*. Only at the early, MCSF-dependent macrophage stage, did we find a significant decrease in the *Tnfrsf11a* encoding RANK in *Sdc3^−/−^* vs WT (Fig.3g), which may explain lower sensitivity of *Sdc3^−/−^* macrophage-lineage cells to RANKL and consequently impaired osteoclastogenesis. The expression of the major transcription factor inducing osteoclastogenesis, *Nfatc1*, increased during osteoclast differentiation, as did the late osteoclast markers *Dcstamp* and *Ctsk* as expected, with no significant differences between genotypes. Interestingly *Sdc3* expression significantly increased in osteoclasts vs MCSF-dependent macrophages, suggesting a role for SDC3 in mature osteoclasts (Fig.3g). Given that during osteoclast formation RANK expression has been shown to be induced by the FZD-mediated non-canonical WNT signalling pathway ROR/JNK/c-JUN^31^ and FZD7 is highly expressed^32^, whereas in *Xenopus* FZD7 associates with SDC4 and RSPO3 in the non-canonical WNT/PCP pathway during endocytosis^16^, we assessed *Fzd7* expression during osteoclastogenesis. Interestingly, again *Fzd7* expression was significantly lower in *Sdc3^−/−^* MCSF-dependent macrophages, but not at the pre-osteoclast or osteoclast stage (Fig.3g). As FZD7 is involved in both canonical and non-canonical WNT-signalling^16,33,34^ and is likely induced by β-catenin itself through the TCF/LEF promoter binding site^35^, taken together our data suggest that SDC3 enhances early stages of osteoclast differentiation via WNT-signalling and plays a role in mature osteoclasts, however the exact mechanism of action of SDC3 in osteoclast lineage cells is currently unclear.

### Low bone volume phenotype in adult *Sdc3^−/−^* mice is due to impaired osteoblast maturation and reduced bone formation

Dynamic histomorphometry of bone formation using calcein double-label quantification in 3 month-old *Sdc3^−/−^* mice revealed a decrease of 35% in BV/TV (*p<0.001*), 21% in the mineral apposition rate (MAR, *p<0.01*), 36% in the mineralising surface per bone surface (MS/BS (*p<0.05*) and 50% in the bone formation rate per bone surface (BFR/BS (*p<0.001*) vs WT (Fig.4a and b, and Supplementary Table 3) in keeping with low bone formation rate in *Sdc3^−/−^* mice. Functional assays confirmed an over 2- and 6-fold reduction in mineralisation in *Sdc3^−/−^* vs WT cultures of osteoblasts differentiated from BMSCs and from BCs respectively (Fig.4c and d). *Sdc3^−/−^* osteoblasts grown out of calvariae also showed significantly reduced mineralisation capacity vs WT (Fig.4c and d). Unsurprisingly, alkaline phosphatase activity was reduced by 70% in *Sdc3^−/−^* vs WT osteoblasts differentiated from BCs *in vitro* (*p<0.001*; Fig.4e), indicating that the low bone formation rate seen in *Sdc3^−/−^* mice on histomorphometry is due to impaired osteoblast function. As the proliferation rate of osteoblasts assessed by the Ki67 assay was higher in *Sdc3^−/−^* compared to WT cultures (9.1±1.7 vs 7.0±1.1, *p<0.05*), taken together these data suggest that SDC3 is important for timely maturation of proliferating osteoblasts and normal bone formation and mineralisation.

**Fig. 4.**
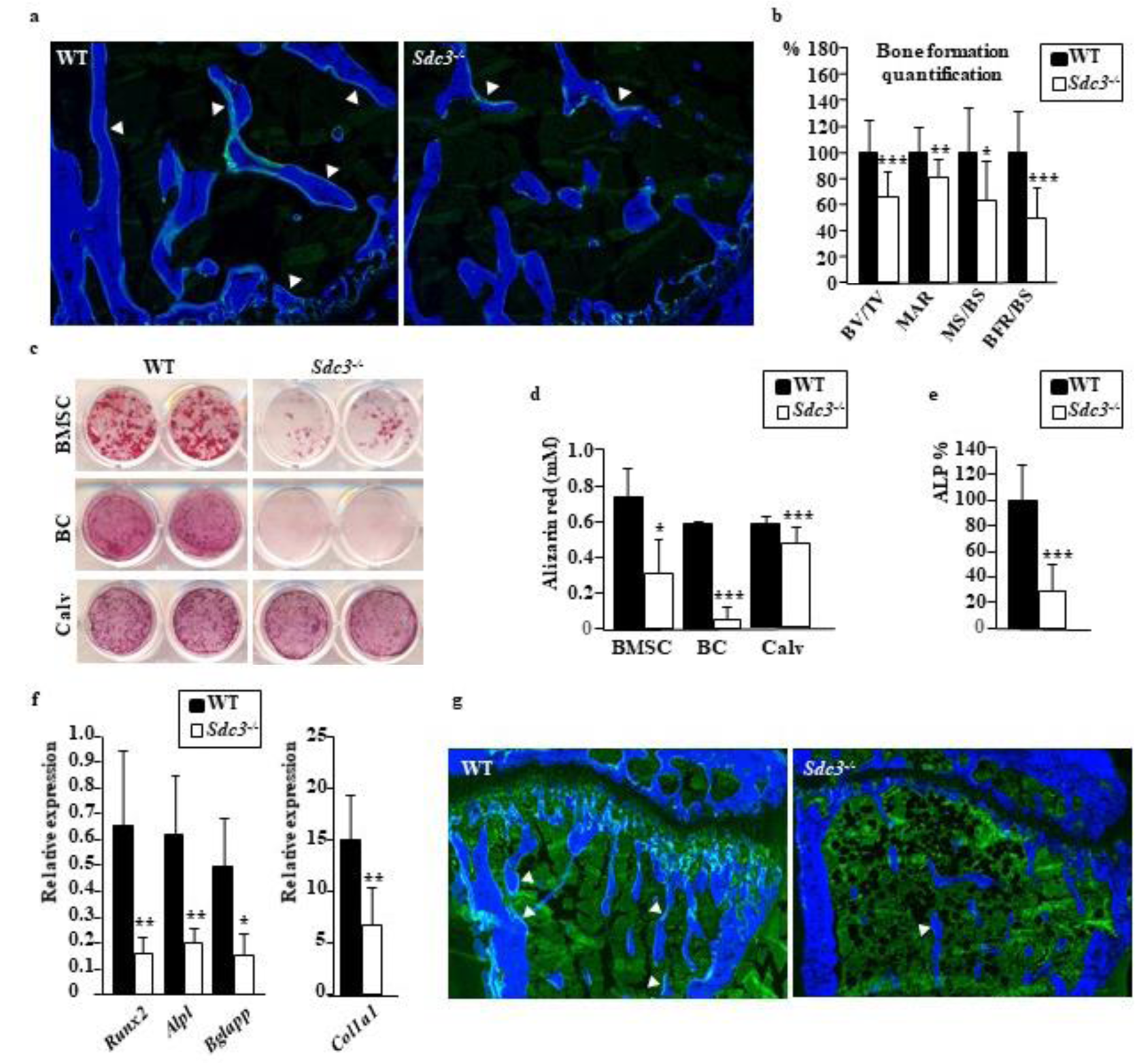
Syndecan-3 enhances bone formation in adult mice. **(a)** Histomorphometry of tibiae from calcein-double labelled 3-month old WT (left panel) and *Sdc3^−/−^* (right panel) mice, bone counterstained with calcein blue. Double label is indicated by arrowheads. **(b)** Quantification of bone formation in calcein-double labelled 3-month old WT (n=12) and *Sdc3^−/−^* (n=12) mice. BV/TV: bone volume/tissue volume, MAR: mineral apposition rate, MS/BS: mineralising surface/bone surface, BFR/BS: bone formation rate/bone surface. **(c)** Photographs of representative WT and *Sdc3^−/−^* osteoblast cultures stained with alizarin red to show mineralisation. In top, middle and bottom row osteoblasts differentiated from bone marrow stromal cells (BMSC), bone chips (BC) and calvariae (Calv) respectively. **(d)** Alizarin red quantification of mineral from cultures of WT and *Sdc3^−/−^* osteoblasts differentiated from BMSC (n=3, 4 replicate wells each), bone chips (BC, n=3, 2 replicates each) and calvariae (Calv, n=3, 2 replicates each) as indicated. **(e)** Alkaline phosphatase (ALP) quantification in WT and *Sdc3^−/−^* osteoblasts grown out from bone chips, data normalised to WT (n=4, 5 replicates each) **(f)** RNA expression of osteoblast marker genes: *Runx2*, *Alpl* (encoding ALP), *Bglap* (encoding osteocalcin) and *Col1a1* relative to *Hmbs assesse*d by qPCR in osteoblasts grown out of bone chips from WT (n=3) and *Sdc3^−/−^* (n=3) mice. **(g)** Histomorphometry of mechanically loaded tibiae from calcein-double labelled 3-month old WT (left panel) and *Sdc3^−/−^* (right panel) mice, bone counterstained with calcein blue. Double label is indicated by arrowheads. Quantification is shown in Table 1. Data in b, d, e and f are shown as mean±SD, * *p<0.05*, ** *p<0.01*, *** *p<0.001*

The functional impairment of *Sdc3^−/−^* osteoblasts was corroborated by the finding of downregulated osteoblastic differentiation and activity genes: *Runx2* by 75% (*p<0.01*), *Alpl* (encoding alkaline phosphatase) by 70% (*p<0.01*), *Bglap* (encoding osteocalcin) by 70% (*p<0.05*) and *Col1a1* by 55% (*p<0.01*) vs WT (Fig.4f). In addition, we quantified *Sdc1*, *Sdc2* and *Sdc4* expression in osteoblasts, to assess whether other syndecans might be increased in an effort to compensate for lack of *Sdc3*, however found no differences between *Sdc3^−/−^* and WT osteoblasts (Supplementary Fig.1).

As bone responds with new bone formation to mechanical loading, we assessed this anabolic response *in vivo*. As expected, in WT mice mechanical loading of the tibia induced an increase in mineralising surface (MS/BS) and in bone formation rate (BFR/BS) of 41% (*p<0.001*) and 48% (*p<0.05*) respectively in the loaded vs non-loaded tibia. However, in *Sdc3^−/−^* mice the anabolic response to mechanical loading was blunted: the increase in MS/BS and BFR/BS in the loaded vs non-loaded tibia was only 12% and 21% respectively (Fig.4h and Table 1), and not statistically significant.

**Table 1.** Anabolic response to mechanical loading analysed by histomorphometry. The right tibia of 10-week-old male WT and Sdc3^−/−^ mice was mechanically loaded for 2 weeks as described, whereas the left tibia was not loaded. Anabolic response was quantified by % change in loaded vs non-loaded tibia and statistical significance calculated using paired Student’s *t* test. Values shown are means±SD, \**p<0.05* and \*\*\**p<0.001*. MAR: mineral apposition rate, MS/BS: mineralising surface / bone surface, BFR/BS: bone formation rate / bone surface.

|  | WT (n=7) |  |  | *Sdc3*<sup>-/-</sup> (n=8) |  |  |
| --- | --- | --- | --- | --- | --- | --- |
|  | Non-loaded | Loaded | % change | Non-loaded | Loaded | % change |
| MAR | 1.97±0.27 | 2.04±0.23 | 4.57±14.66 | 1.60±0.28 | 1.65±0.26 | 3.64±17.10 |
| MS/BS | 27.53±8.49 | 38.37±10.93 | 40.72±17.83*** | 16.10±5.22 | 18.14±7.43 | 11.65±32.48 |
| BFR/BS | 0.54±0.18 | 0.80±0.30 | 48.77±29.19* | 0.26±0.09 | 0.31±0.15 | 20.84±46.76 |

### Bone marrow adiposity is increased in *Sdc3^−/−^* mice

As *Sdc3* deletion leads to a bone phenotype mimicking premature osteoporosis, which in humans is typically associated with increased bone marrow adipose tissue (BMAT), we quantified BMAT, seen as empty spaces in the bone marrow on the Goldner’s Trichrome stain, after the void areas had been confirmed to contain adipocytes on perilipin staining (Fig.5a). Histomorphometry analysis of tibial and femoral bones from 3-month old mice revealed a 60-fold increase in bone marrow adipocyte number and area in *Sdc3^−/−^* compared to WT (*p<0.001*, Fig.5b and c). *In vitro*, a two-fold increase in adipocyte formation was observed in *Sdc3^−/−^* vs WT BMSCs cultured in adipo-osteoblastogenesis conditions (Fig.5d-f) and adipocytes formed 1-2 days earlier in *Sdc3^−/−^* than WT cultures. Thus, deletion of *Sdc3* leads to a premature osteoporosis-like phenotype characterised by not only low bone volume and fragility, but also increased BMAT, possibly due to a preferential switch from osteoblastogenesis to adipogenesis at a progenitor level.

**Fig. 5.**
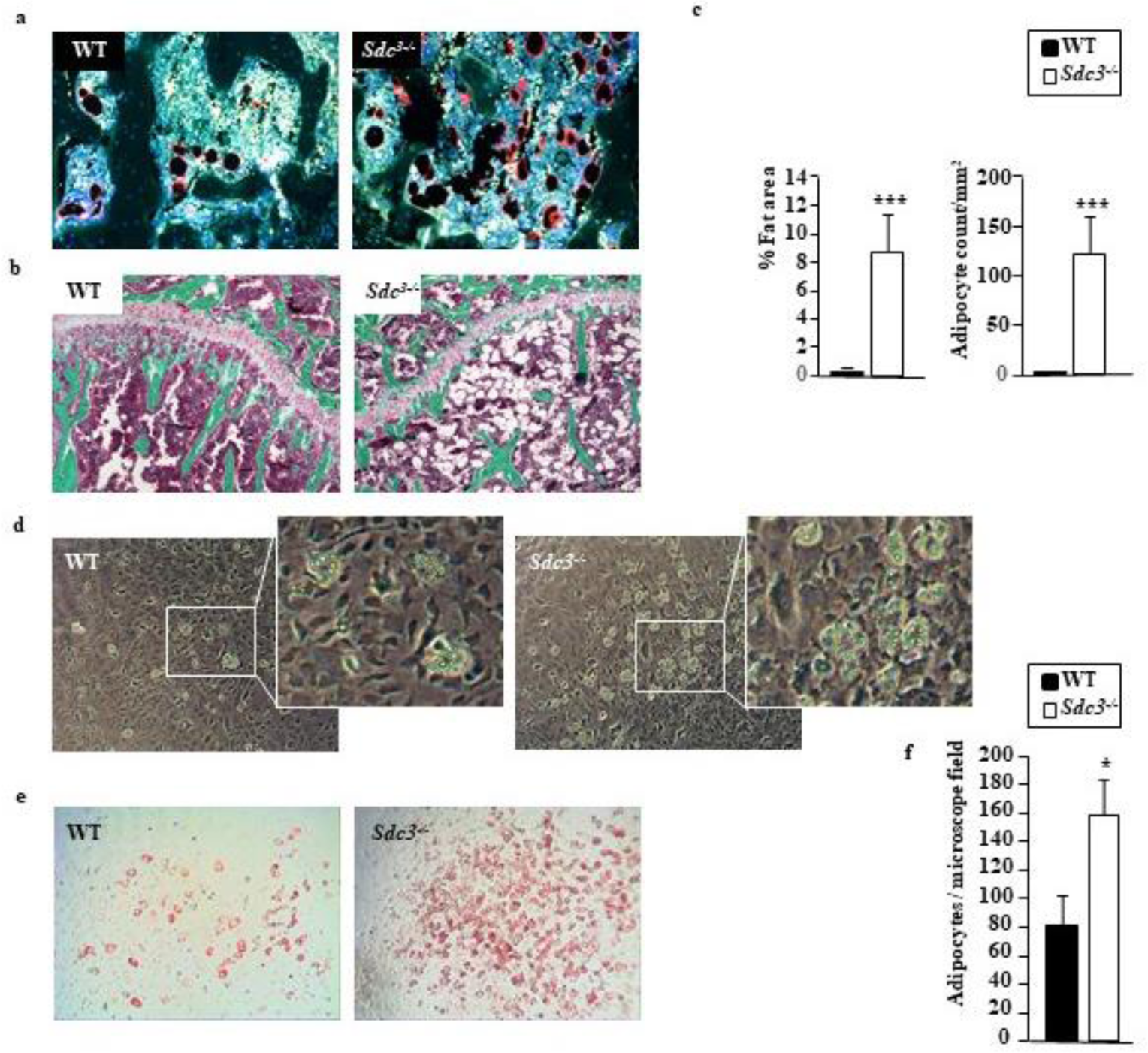
Syndecan-3 deletion leads to an increase in bone marrow adipose tissue (BMAT) **(a)** Perilipin stain of bone marrow adipocytes (seen as black ovals with a red rim) in proximal tibia from 3-month old WT (left panel) and *Sdc3^−/−^* (right panel) mice. **(b)** Goldners trichrome stain of proximal tibia from 3-month old WT (left panel) and *Sdc3^−/−^* (right panel) mice, showing oval voids corresponding to adipocytes within the bone marrow. **(c)** Quantification of BMAT area and adipocyte count in WT (n=6) and *Sdc3^−/−^* (n=4) mice on histology. **(d)** Representative images of WT (left panel) and *Sdc3^−/−^* (right panel) BMSCs grown in adipo-osteogenic conditions for 12 days, fixed and visualised by phase-contrast microscopy at 10x magnification. White rectangles delineate clusters of adipocytes magnified in the smaller panels to the right. **(e)** Representative images of BMSC cultures described in d stained with Oil-red-O to visualise adipocytes. Bright field microscopy, magnification 10x. **(f)** Quantification of adipocytes from WT (n=3) and *Sdc3^−/−^* (n=3) BMSC cultures described in d and e. Data in c and f are mean±SD, * *p<0.05*, *** *p<0.001*

### Syndecan-3 enhances canonical WNT signalling pathway in osteoblasts

Given that a common signalling pathway enhancing osteoblastogenesis, mediating anabolic response to mechanical loading and in cartilage critical for normal growth plate development is WNT-mediated^4^,^36^, we hypothesised that SDC3 enhances WNT signalling.

To assess the effect of *Sdc3* deletion on canonical WNT-signalling, osteoblasts were stimulated with WNT3a, which has been shown to enhance osteoblastogenesis^37,38^, and the main β-catenin target gene *Axin2* quantified by qPCR. In *Sdc3^−/−^* osteoblast cultures there was a significant reduction in *Axin2* expression compared to WT controls regardless of whether the osteoblasts were grown out of BCs or differentiated from BMSCs, which was mostly apparent after stimulation with WNT3a (Fig.6a) and in keeping with decreased canonical WNT-signalling in the absence of SDC3.

**Fig. 6.**
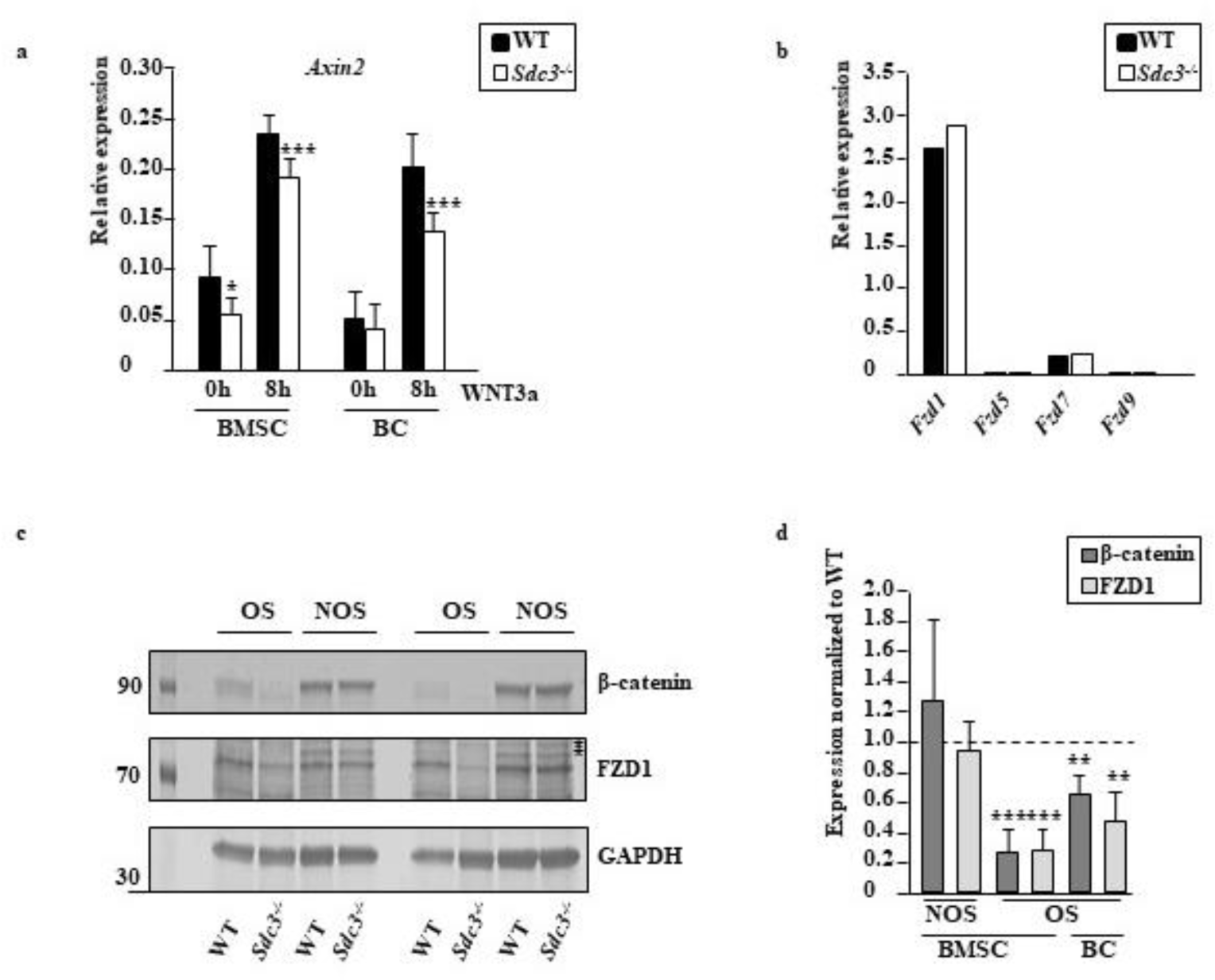
Syndecan-3 deletion enhances Frizzled 1 degradation and impairs canonical WNT signalling in osteoblasts. **(a)** Osteoblasts were differentiated from bone marrow mesenchymal stromal cells (BMSC) or bone chips (BC) from *Sdc3^−/−^* (n=3) or WT (n=3) mice and grown in osteogenic conditions. 16 hours prior to stimulation with WNT3a FCS was replaced with TCM and *Axin2* expression relative to *Hmbs* was quantified by qPCR in triplicates. Values shown are means±SD. \**p<0.05*, \*\**p<0.01*, \*\*\**p<0.001*. **(b)** Osteoblasts were differentiated from BC and grown until confluence. *Fzd1, 5, 7 and 9* expression relative to *Hmbs* was quantified by qPCR in triplicates (n=1). **(c)** Osteoblasts were differentiated from *Sdc3^−/−^* (n=5) and WT (n=5) BMSCs and grown in non-osteogenic (NOS) or osteogenic (OS) conditions. Cell lysates were immunoblotted for β-catenin, FZD1 and GAPDH. Western blot shows 2 representative experiments out of 5. Asterisks indicate non-specific bands. **(d)** Immunoblot quantification of β-catenin and FZD1 protein expression in *Sdc3*^−/−^ osteoblasts differentiated from *Sdc3^−/−^* (n=5) and WT (n=5) BMSCs (as per c) or BCs (*Sdc3^−/−^* n=5, WT n=5) grown in NOS or OS conditions until confluent. All β-catenin and FZD1 bands were normalised to GAPDH and *Sdc3^−/−^* data were normalised to respective WT. Data are shown as means±SD. \*\**p<0.01*, \*\*\**p<0.001*

Thus, we further assessed the main canonical WNT-signalling pathway components at mRNA level and found no difference in expression of *Lrp5* or *6*, *Lgr4-6*, *Znrf3* and *Rnf43*, in *Sdc3^−/−^* vs WT osteoblasts (data not shown). From the *Fzds* involved in canonical WNT signalling (i.e. *Fzd1*, *5*, *7* and *9*), *Fzd1* expression in osteoblasts was highest (**Fig.6b**) in keeping with previous reports^39^. As FZD1 enhances osteoblast-mediated mineralisation^40^, we quantified FZD1 and β-catenin at protein level and found both reduced by 72% and 73% respectively in *Sdc3^−/−^* vs WT (*p<0.001*) in BMSC-derived osteoblasts grown in osteogenic conditions (Fig.6c and d). In osteoblasts derived from bone-chips, FZD1 and β-catenin were reduced by 52% (*p<0.*01) and 34% (*p<0.*01) respectively in *Sdc3^−/−^* vs WT (Fig.6c and d). In contrast, β-catenin levels were considerably higher in BMSCs cultured in non-osteogenic conditions, and there was no difference in either FZD1 or β-catenin levels between *Sdc3^−/−^* and WT in these cells. Taken together, these data suggest that in osteoblasts SDC3 enhances the canonical WNT-signalling pathway by stabilising FZD1.

## DISCUSSION

Here we have identified a novel anabolic role for SDC3 in bone. Deletion of *Sdc3* leads to low bone volume, increased bone fragility and increased bone marrow fat in young adult mice, a premature osteoporosis-like phenotype.

Interestingly, SDC3 differentially affects pre- and post-natal long bone growth. *Sdc3* deletion resulted in increased long bone elongation *in utero*, in keeping with the known restrictive role of SDC3 at two stages of skeletal development. Firstly, during the condensation stage of early pre-cartilaginous skeletogenesis SDC3 restricts the size of mesenchymal cell aggregates and promotes the differentiation step^18,41,42^. Secondly, during limb elongation SDC3 is thought to set boundaries at the metaphyseal and epiphyseal perichondrium and periosteum^41,43^. Thus, deletion of *Sdc3* would allow for a relative expansion of the cartilaginous skeleton, explaining the longer bones seen early postnatally in *Sdc3^−/−^* mice. Moreover, it has been previously shown that SDC3 regulates proliferation and maturation of chondrocytes in the growth plate during embryonic long bone development^20,44,45^ and restricts chondrogenic differentiation *in vitro*^30^. Our data indicate that SDC3’s regulatory role extends into adulthood as SDC3 deletion leads at 3 months of age to a thin growth plate with disorganised columns of chondrocytes within the proliferating zone, which is associated with mild, but significant long bone growth stunting compared to the reverse seen *in utero*. This growth plate abnormality may be due to defective chondrogenesis at stem cell level in the growth plate resting zone^46,47^, and/or at the proliferation and differentiation stages of chondrocytes^19,20^, in which extracellular matrix, with which transmembrane HSPGs interact, is thought to play a role. Further investigations are in progress to assess the exact role of SDC3 in the regulation of chondrocytes within the growth plate and in longitudinal bone growth.

Significantly, the relatively large total long bone volume of *Sdc3^−/−^* pups is not only not maintained into adulthood but reversed already at 2 weeks of age, whereas the trabecular bone volume (BV/TV) becomes significantly lower at 2 months and remains low. Whilst the shortening of the long bones in *Sdc3^−/−^* adult mice reflects a thinner and morphologically abnormal growth plate, the low bone volume phenotype is unexpected. However, a similar long-bone phenotype has been reported in young adult mice deficient in the heparin-binding growth molecule (HB-GAM, also known as osteoblast stimulating factor 1 [OSF1] and pleiotrophin), expressed at the growth plate and around osteocytes in bone^48,49^. Thus, our findings indicate that SDC3, like HB-GAM, mediates an osteogenic effect in response to mechanical loading, given that SDC3 likely serves as receptor for HB-GAM on osteoblasts, as has been previously shown in neurons^50^.

Although osteoclastogenesis is decreased in *Sdc3^−/−^* mice (like in *HB-GAM^−/−^*), the relative scarcity of osteoclasts is likely offset by increased osteoclast-mediated bone resorption, which may be contributory to the low bone volume phenotype. The latter finding contrasts with the lack of effect of HB-GAM on osteoclastic bone resorption^48^. Our observation of a gradual increase in SDC3 expression during normal osteoclast differentiation suggests its importance for osteoclast maturation and/or function. Indeed, reduced expression of *Tnfrsf11a* encoding RANK and *Fzd7* in MCSF-dependent *Sdc3^−/−^* macrophages, is reflected by reduced sensitivity to RANKL and suggests that SDC3 may enhance pathways targeting *Fzd7*. These include upstream from *Fzd7* the canonical β-catenin mediated WNT^35,51^ and the non-canonical WNT signalling mediated by FZD via RAC/JNK/c-Jun inducing expression of RANK^31^. On the other hand, it is tempting to speculate that SDC3 may associate with FZD7 during cell polarisation, not unlike SDC4 during the planar cell polarity (PCP) process in Xenopus, which also leads to activation of JNK^16^. Furthermore, in human breast epithelial cells FZD7 is induced by NOTCH signalling^52^, which is known to crosstalk with SDC3 during myogenesis^53^, thus further research is currently ongoing exploring the SDC3-NOTCH crosstalk in bone. Our findings are not explained by the lack of the SDC3 ectodomain in the light of Kim’s and colleagues recent report of a suppressive effect of SDC1-4 ectodomains on osteoclastogenesis and of SDC1, 2 and 4 (but not SDC3) ectodomains on osteoclast mediated bone resorption^54^.

The *Sdc3^−/−^* mouse low bone volume phenotype primarily reflects impaired osteoblastic bone formation, which is associated with increased bone marrow adiposity, suggesting either a preferential switch at the mesenchymal progenitor level from osteoblastogenesis to adipogenesis, or potential de-differentiation of osteoblasts towards adipocyte lineage. It is unlikely that systemic factors play a significant role in this aspect of the phenotype, as at a cellular level there is increased adipocyte formation seen in *Sdc3^−/−^* BMSCs vs WT *in vitro*. Indeed, the delay in *Sdc3^−/−^* osteoblast differentiation suggests a degree of immaturity of differentiated *Sdc3^−/−^* osteoblasts, which display impaired ability to not only form, but also fully mineralise newly formed bone. The blunted anabolic response of bone to mechanical loading in *Sdc3^−/−^* mice is not due to muscle dysfunction as deletion of *Sdc3* in fact improves muscle homeostasis, regeneration and ageing, in both healthy and dystrophic mice^55^. Moreover, *Sdc3^−/−^* mice are no different from WT in regard to endurance training performance^55^.

As the key pathways regulating skeletal development, inducing osteoblastogenesis^4^, inhibiting adipogenesis^56,57^, and in cartilage inducing chondrocyte hypertrophy and regulating the morphology of the growth plate^36,58^ involve WNTs^4^, we focussed our investigations on the canonical WNT signalling. To our knowledge, involvement of SDC3 in WNT signalling has not been reported previously, although SDC1, 2 and 4 are co-receptors in WNT signalling^17^. Given the severity of the *Sdc3^−/−^* mouse bone phenotype, clearly SDC1, 2 and 4 are unable to compensate for lack of SDC3, indicating its specific role in the acquisition of bone volume and maintenance of normal bone homeostasis in adult mice.

We consistently found decreased expression of *Axin2* in *Sdc3^−/−^* osteoblasts indicating impaired β-catenin signalling. Out of the *Fzds* classically involved in canonical WNT signalling, i.e. *Fzd1*, *5*, *7* and *9*^17^, we found *Fzd1* expression in osteoblasts to be highest, as reported previosuly^39^. Given that FZD1 not only mediates osteoblast differentiation, but also bone mineralization^40^ and that *Sdc3^−/−^* osteoblasts show impairment in both these functions, our findings of decreased FZD1 and β-catenin in *Sdc3^−/−^* osteoblasts may explain the *Sdc3^−/−^* phenotype. Furthermore, as *Fzd1* RNA expression is not affected in *Sdc3^−/−^* osteoblasts, whereas at the protein level FZD1 is significantly decreased in committed *Sdc3^−/−^* osteoblasts, it is plausible that in the absence of SDC3, FZD1 undergoes rapid degradation, leading to decreased β-catenin signalling. Thus, we propose that in osteoblasts SDC3 enhances canonical WNT signalling and likely serves as co-receptor which stabilises FZD1, however, the exact molecular mechanism of this interaction remains to be explored. Interestingly, β-catenin levels were considerably higher in BMSCs grown under non-osteogenic conditions than in those cultured for 7 days in osteogenic conditions, findings similar to previously reported in a human pre-osteoblast SV-HFO cell line^39^. This may indicate that during osteoblast differentiation β-catenin levels, already low, are more critical and tightly controlled. Thus, any changes in FZD1 levels may have a relatively more pronounced effect on β-catenin signalling.

Although our findings indicate that SDC3 enhances canonical WNT signalling in the osteoblast lineage and both canonical and non-canonical WNT signalling in osteoclast lineage, other pathways remain to be investigated. These, in addition to the remaining WNT signalling pathways, include FGF, BMP and NOTCH, crosstalk of which orchestrates osteoblast differentiation^59^, given that SDCs have been shown to partake in all these pathways^60^. Importantly, interaction of SDC3 with HB-GAM has been shown to regulate osteoblast recruitment to sites of increased bone formation^49^ and osteocytic HB-GAM expression is upregulated after mechanical loading^48^, thus investigation of this pathway in the *Sdc3^−/−^* mouse is ongoing.

In summary, SDC3 differentially affects pre- and postnatal bone development. SDC3 deletion leads to a low bone volume phenotype associated with high BMAT in young adult mice. The anabolic effect of SDC3 on bone is mediated through canonical WNT signalling in osteoblasts, likely through stabilisation of FZD1. Increased bone resorption, but decreased osteoclastogenesis in *Sdc3^−/−^* mouse, likely contributes to the low bone volume phenotype. Thus, SDC3 is a novel therapeutic target for anabolic drug development for treatment of osteoporosis.

## ACKNOWLEDGEMENTS

We would like to thank Andy Houghton and Euan Owen, Institute of Ageing and Chronic Disease, University of Liverpool, for their help with animal work. This project was financially supported by the Medical Research Council (grant MR/R002819/1) and funding from the Institute of Ageing and Chronic Disease, University of Liverpool.

## AUTHOR CONTRIBUTIONS

A.D. and R.J.H. designed the experiments with input from A.B., A.P.^2,3^, G.B.G. and B.P.; F.M.J.S.B., A.B., A.P.^1^, G.C. and C.S. performed the experiments and acquired the data; F.M.J.S.B., A.B., A.P. ^2,3^, B.P., M.S.M., G.B.G., R.J.H. and A.D. analysed the data; A.D. and R.J.H. wrote the manuscript, which was critically revised by all co-authors.

## COMPETING INTERESTS

The authors declare no competing interests.

---

**Supplementary Fig. 1.**
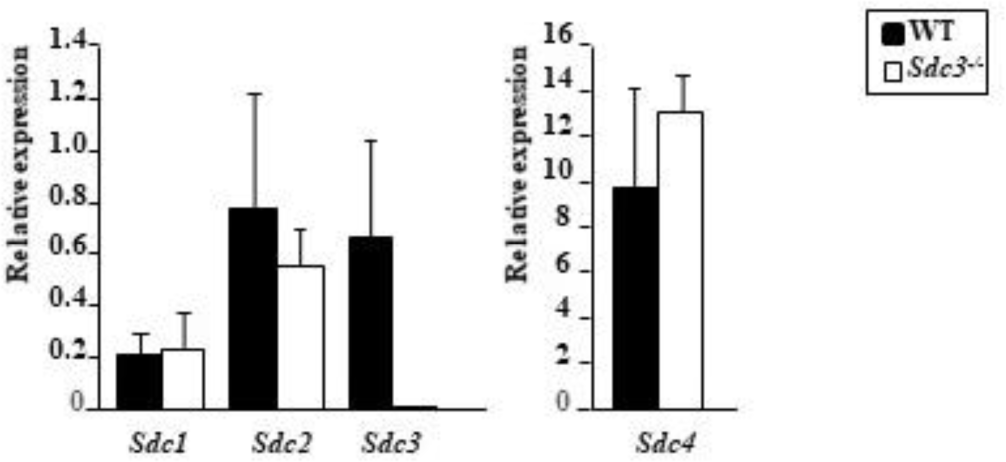
Syndecan expression in osteoblasts. RNA expression of *Sdc1*, *Sdc2*, *Sdc3* and *Sdc4* relative to *Hmbs assesse*d by qPCR in osteoblasts grown out of bone chips from WT (n=3) and *Sdc3^−/−^* (n=3) mice. Data are shown as mean±SD.

**Supplementary Table 1.**
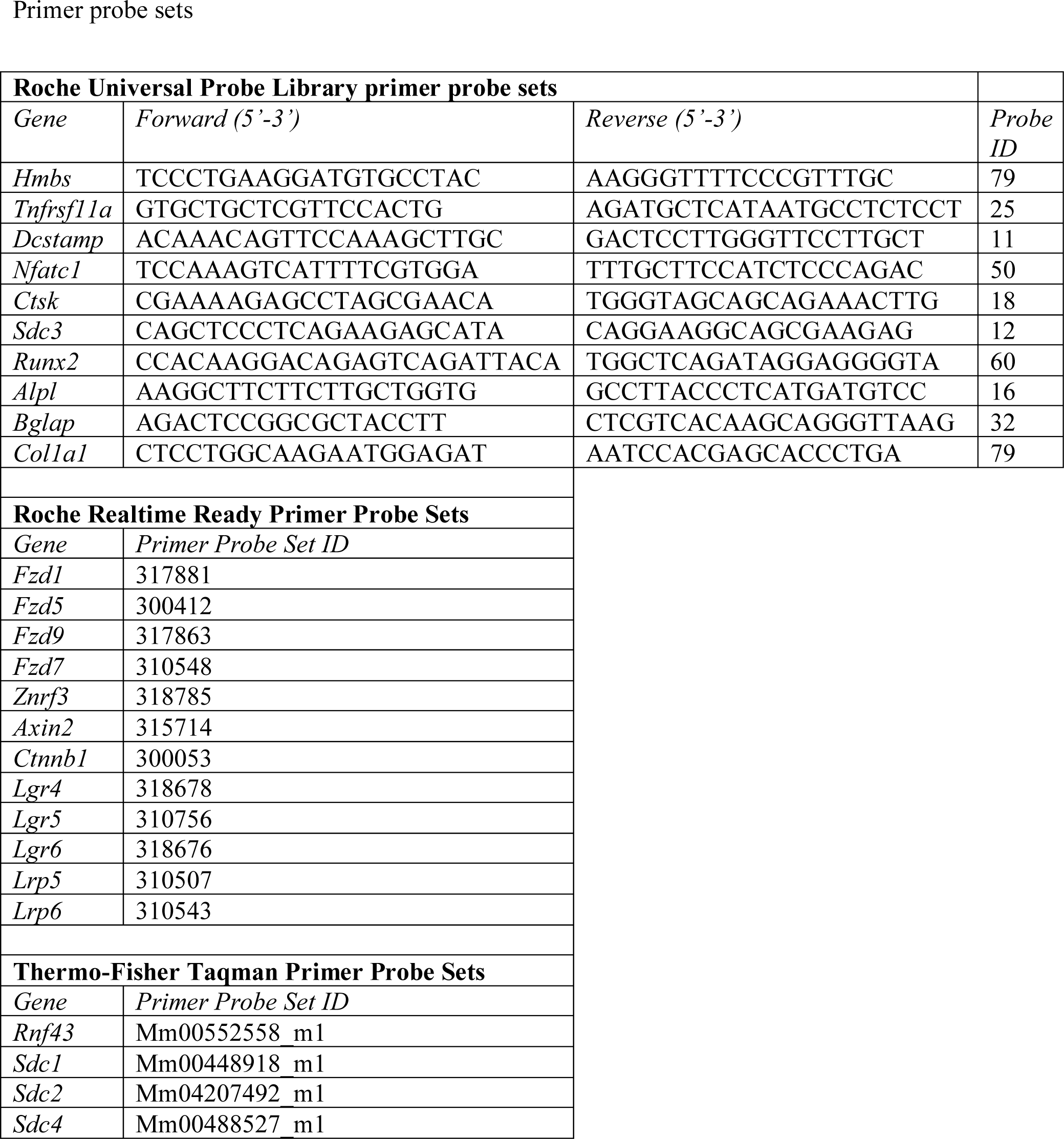

**Supplementary Table 2a.**
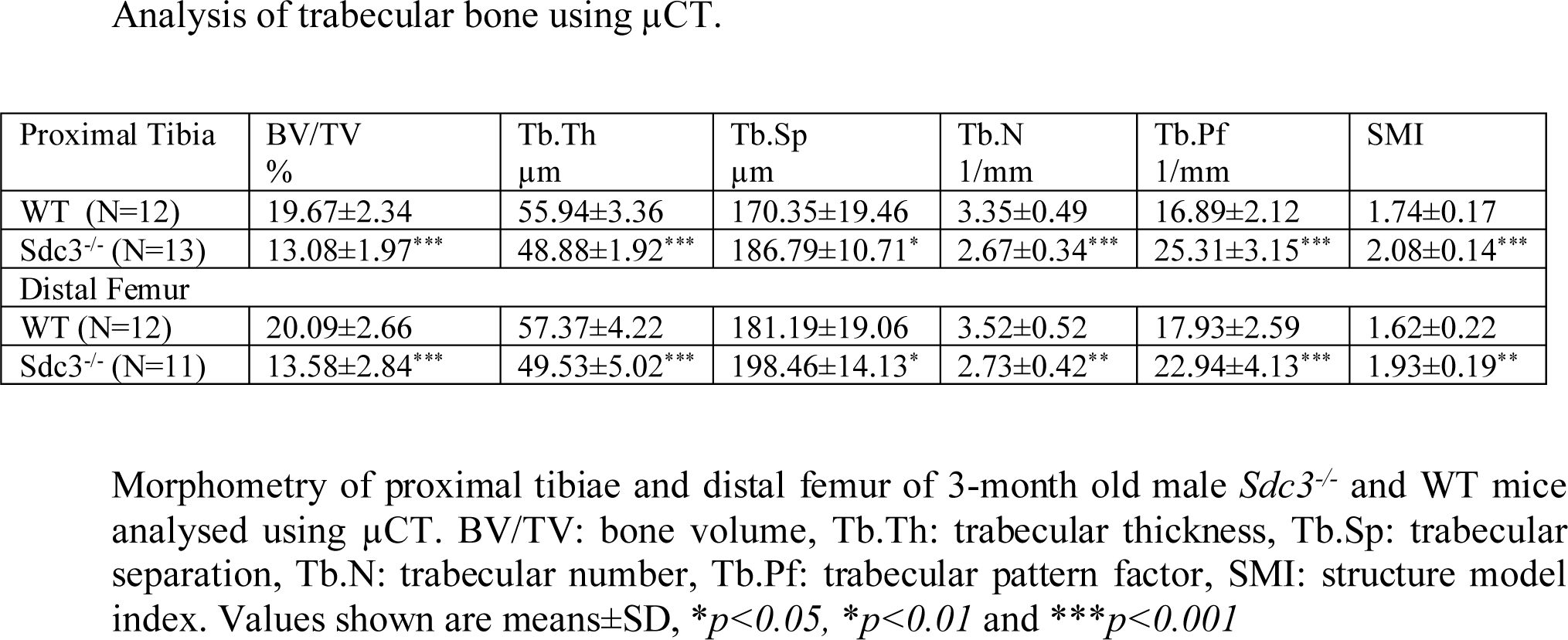

**Supplementary Table 2b.** Analysis of cortical bone using μCT.

| Tibia | Cort.Th<br>μm | E.Pm<br>mm | P.Pm<br>mm | Polar MMI<br>mm<sup>4</sup> |
| --- | --- | --- | --- | --- |
| WT (N=12) | 259.82±15.56 | 2.34±0.15 | 4.24±0.52 | 0.20±0.07 |
| Sdc3<sup>-/-</sup> (N=10) | 237.30±9.7** | 2.23±0.08 | 3.86±0.10* | 0.13±0.05* |
| Femur |  |  |  |  |
| WT (N=9) | 224.80±14.53 | 4.76±0.32 | 5.89±0.21 | 0.56±0.10 |
| Sdc3<sup>-/-</sup> (N=11) | 201.15±13.64*** | 4.21±0.23*** | 5.54±0.11** | 0.44±0.07** |

Morphometry of tibial and femoral cortex of 3-month old male Sdc3<sup>-/-</sup> and WT mice analysed using μCT. C.Th: cortical thickness, E.Pm: endosteal perimeter, P.Pm: periosteal perimeter, MMI: mean moment of inertia. Values shown are means ± SD, \* $p$ <0.05, \*\* $p$ <0.01 and \*\*\* $p$ <0.001

**Supplementary Table 3.**
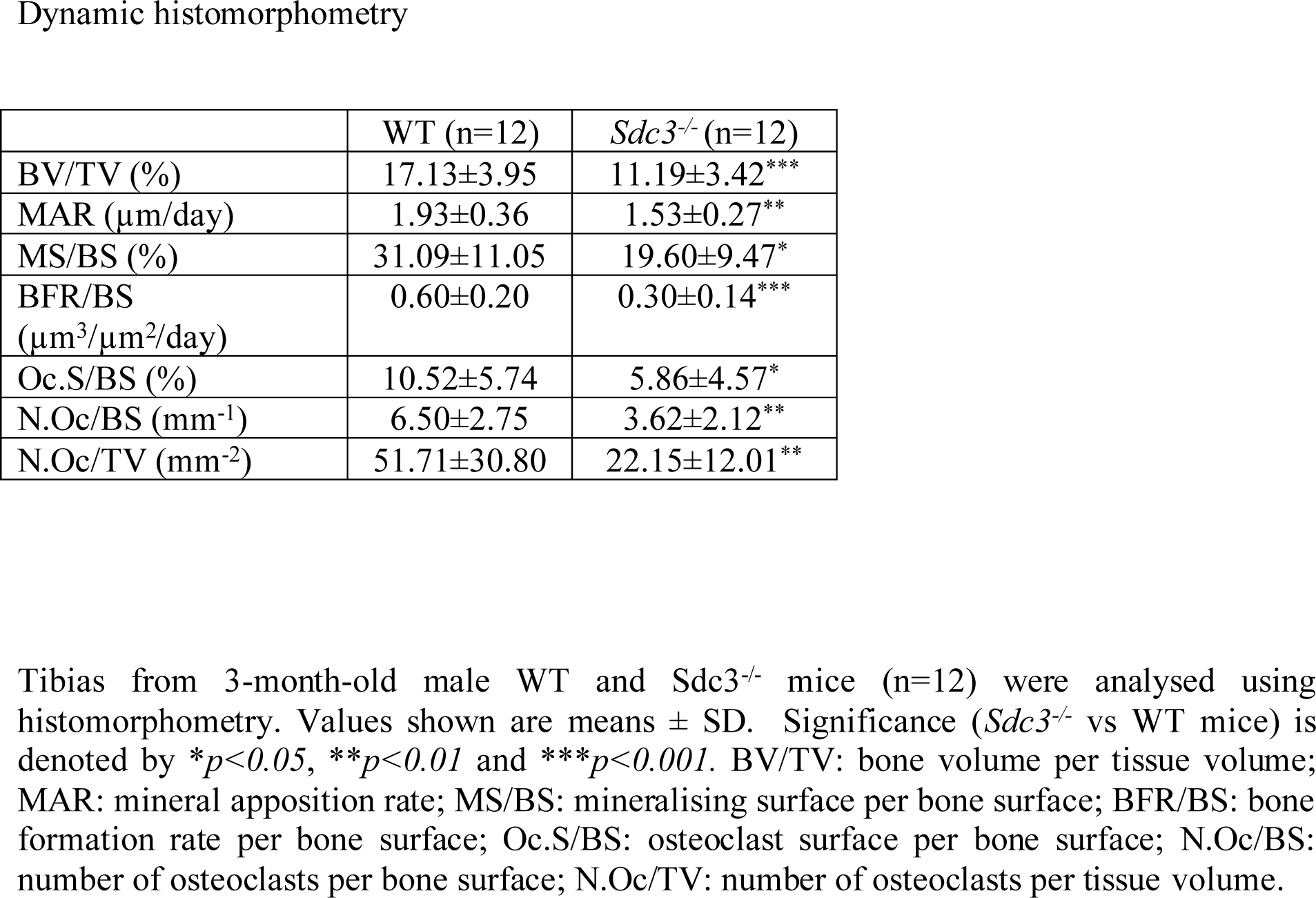

